# Comparative evaluation of reference genomes and cell-type annotation frameworks for single-nucleus transcriptomic analysis in apple

**DOI:** 10.64898/2025.12.02.691976

**Authors:** Shupei Hu, Yiqi Liu, Yujie Zhao, Bo Li, Yawen Shen, Yoshiharu Mimata, Kunxi Zhang, Pengbo Hao, Jiangli Shi, Zhiwei Luo, Wenxiu Ye, Xianbo Zheng, Jian Jiao

**Affiliations:** College of Horticulture, Henan Agricultural University, Zhengzhou 450046, China; Peking University Institute of Advanced Agricultural Sciences, Shandong Laboratory of Advanced Agricultural Sciences at Weifang, Weifang, Shandong 261325, China; Plant and Food Research, Bioeconomy Science Institute, Auckland 1025, New Zealand; Zhongyuan Research Center, Chinese Academy of Agricultural Sciences, Xinxiang 453500, Henan, China

**Keywords:** snRNA-seq, *Malus domestica*, Reference genome, Cell type annotation, Pseudotime analysis

## Abstract

Single-nucleus RNA sequencing (snRNA-seq) provides a powerful tool to profile cell-type-specific transcriptional programs in woody fruit trees. Yet its application in *Malus domestica* has been limited by heterogeneous reference genomes and the lack of validated marker genes. Here, we first established a comprehensive set of marker genes corresponding to 29 plant cell types and identified their orthologs in apple. Using these resources, we systematically evaluated six publicly available apple genome assemblies as references for snRNA-seq analysis. Our results demonstrate that reference genome selection has a major impact on mapping performance and downstream cell type annotation, with the GDDH13 assembly consistently producing the highest transcript counts per nucleus and the greatest number of identifiable transcriptional clusters (15 clusters representing 11 distinct cell types across 35,557 high-quality nuclei). To achieve robust cell type annotation, we integrated classical marker gene-based annotation with two cross-species computational frameworks — Orthologous Marker Gene Groups (OMGs) and XSpeciesSpanner. These three approaches enabled confident identification of 11 major cell types in apple seedlings and provided cross-validation for ambiguous or previously unannotated clusters. Pseudotime analysis reconstructed the developmental trajectories of both procambial and stomatal lineages, revealing their progressive transcriptional transitions and enabling the delineation of key molecular programs underlying vascular differentiation and early guard-cell formation. Together, this work provides a reference-quality single-nucleus transcriptomic atlas for apple and underlined the critical impact of genome choice and multi-method cell-type annotation on single-cell studies in non-model crops.

## 1 Introduction

Apple (Malus domestica) ranks among the most widely cultivated and economically important fruit crops worldwide, with global annual production surpassing 85 million tons [1]. It has also emerged as an ideal species for woody fruit tree research, owing to its well-annotated genome, extensive genetic diversity, and the availability of modern biotechnological tools [2–4]. Despite these advances, our understanding of cell-type-specific gene expression patterns in apple remains limited, hindering progress in elucidating cellular functions and developmental processes in this crop [5].

Single-cell RNA sequencing (scRNA-seq) has reshaped plant biology by making it possible to examine gene expression at true single-cell resolution [6]. Over the past few years, researchers have extended its use to a broad range of plant tissues, such as roots, shoot apices, leaves, stems, and inflorescences across multiple species [7]. This technology has transformed developmental and functional studies in model plants, revealing novel cell populations, developmental trajectories, and regulatory networks [8]. However, accurate cell type annotation remains a major challenge in plants, particularly in non-model species where genome resources vary in quality and completeness. Moreover, the limited availability of well-validated marker gene sets is also a major obstacle for snRNA-seq studies [9]. These limitations often lead to ambiguous or inconsistent annotations [10]. Therefore, establishing robust strategies that integrate optimized reference genomes and conserved marker identification is essential for achieving reliable cell-type identity in woody fruit crops such as apple.

Conventional cell-type annotation strategies in non-model plants often borrow marker genes from well-studied species such as *Arabidopsis*. However, gene duplication, subfunctionalization, and lineage-specific expression divergence can lead to inaccurate or ambiguous cell identity assignments [11]. Furthermore, the absence of standardized cross-species annotation frameworks limits our ability to compare cell types and to identify conserved developmental programs across plant lineages [12,13]. To address these limitations, a recent approach termed Orthologous Marker Gene Groups (OMGs) was proposed [9]. Rather than relying on one-to-one ortholog matches, OMGs assemble orthologous marker genes into groups and statistically assess their enrichment across cell clusters, allowing cross-species annotation without the need to merge full transcriptomic datasets [9]. OMGs accommodate complex gene family relationships and thus provide a scalable, flexible framework for annotating cell identities in non-model plants [11]. In parallel, the XSpeciesSpanner framework extends cross-species annotation by quantitatively scoring the similarity between query clusters and well-annotated reference atlases based on marker gene expression patterns [14], offering an independent and complementary line of evidence for cell identity assignment in non-model systems. Yet, their application in species such as apple has not been systematically evaluated, particularly regarding annotation reliability.

In this study, we identified the reference genome that provides more reliable support for downstream analyses of apple snRNA-seq data through on systematic evaluation. Leveraging this optimized foundation, we then combined canonical marker-based annotation with the OMG and XSpeciesSpanner frameworks to generate a high-confidence cell type map of young apple seedlings at single-nucleus resolution. By integrating these complementary approaches, we systematically defined apple cell type–specific marker genes, evaluated their conservation relative to model species, and cross-validated cell identity assignments across methods. Building this curated atlas, we further reconstructed the bifurcating differentiation trajectories of procambial cells toward phloem parenchyma and xylem, and delineated the early stomatal-lineage progression from *SPCH*-expressing precursors to *FAMA*-positive guard cells. Together, this integrated strategy provides a robust paradigm for cell type identification and developmental trajectory analysis in woody fruit crops.

## 2. Materials and Methods

### 2.1 Plant material and Nuclei isolation

Seeds of Malus domestica ‘Royal Gala’ were surface-sterilized with 75% ethanol for 1 min, followed by 2% (v/v) sodium hypochlorite for 15 min, and rinsed five times with sterile distilled water and germinated under controlled growth conditions (24 ± 2°C, 16-h light/8-h dark photoperiod, 100 μmol m^−2^ s^−1^ light intensity, 60% relative humidity). Once the seedlings developed four true leaves, the entire seedlings, including roots, hypocotyls, and shoot tissues, were harvested. To preserve nuclear integrity and minimize transcriptional artifacts, tissue collection was performed rapidly. Fresh seedling tissues (∼100 mg) were immediately flash-frozen in liquid nitrogen and stored at -80℃ until nuclei extraction.

Nuclei isolation was performed following the manufacturer’s instructions of the DNBelab C Single-Cell RNA Library Preparation kit (MGI, Shenzhen, China) with minor modifications. Briefly, frozen tissues were homogenized in ice-cold Homogenization Buffer, and nuclei were subsequently washed in Wash Buffer and resuspended in Blocking Buffer supplied with the kit. Nuclei integrity (>90%) and viability (>85%) were confirmed by microscopy and trypan blue staining before library preparation.

### 2.2 snRNA-seq library construction and sequencing

Single-nucleus RNA-seq libraries were constructed using the DNBelab C Series High-throughput Single-Cell RNA Library Preparation kit (MGI, Shenzhen, China), following the manufacturer’s instructions [15]. The single-nucleus suspension was processed for droplet encapsulation, emulsion breakage, and bead collection to capture mRNAs. This was followed by reverse transcription, cDNA amplification, and purification. The resulting cDNA was sheared into short fragments (300–500 bp), and indexed sequencing libraries were prepared. Library quality was quantified using the Qubit ssDNA Assay Kit (Thermo Fisher Scientific) and the Agilent 2100 Bioanalyzer.

The libraries were sequenced on the DNBSEQ-T7 platform (MGI Tech Co., Ltd.) in paired-end mode. The sequencing strategy generated a 30-bp Read 1 (containing the 10-bp cell barcode 1, 10-bp cell barcode 2, and 10-bp unique molecular identifier [UMI]), a 100-bp Read 2 (containing the genomic insert), and a 10-bp sample index read [16].

### 2.3 snRNA-seq data processing

Raw sequencing reads generated from the DNBSEQ platform were first processed to remove low-quality reads and adaptor sequences using fastp (v.0.19.7) [17]. The clean reads were then aligned to six different apple genome assemblies—GDDH13 v1.1, Fuji T2T, GDT2T, Gala, Granny Smith, and HFTH1—using STAR (v2.7.11b) [18] (Supplementary Table 1). To accommodate the specific barcode structure of the DNBelab C library, read alignment and gene-barcode matrix generation were performed with optimized parameters (Supplementary Table 2).

The resulting filtered gene-barcode matrices were imported into R (v4.3.0; R Core Team, 2023) and processed using the Seurat package (v5.3.0) [19]. Quality control was performed by excluding nuclei with fewer than 200 detected genes and filtering out genes expressed in fewer than three nuclei. Highly variable genes were identified using the FindVariableFeatures function, followed by data scaling using ScaleData. Principal component analysis (PCA) was performed for dimensionality reduction, and the top 30 principal components were selected for downstream analysis. Cell clustering was carried out using the Louvain algorithm (via FindNeighbors and FindClusters) with a resolution parameter of 0.5. Potential doublets were identified and removed using DoubletFinder [20]. Finally, the clusters were visualized using Uniform Manifold Approximation and Projection (UMAP)

### 2.4 Marker gene-based cell type annotation

To assign cell types to the identified clusters, we compiled a list of established marker genes based on previously published gene function studies and recent single-cell transcriptomic analyses (Supplementary Table 3). To identify the corresponding homologs in apple, we retrieved the protein sequences of these reported marker genes and performed BLASTP alignments against the predicted proteomes of each apple reference genome [21]. The top hits with significant sequence similarity were designated as the apple orthologs for downstream annotation (E-value < 1e^-5^). We then manually assigned each cluster a putative identity by cross-referencing its cluster-enriched genes with these identified markers. For each cell, the average normalized expression of the marker genes was calculated, and the resulting expression profiles guided the cluster-to-cell type assignments. In addition to DotPlots, FeaturePlots were used to visualize the expression patterns of specific marker genes across clusters. This multifaceted approach allowed comprehensive validation of marker gene expression and provided further support for the accuracy of our cell type annotations.

### 2.5 OMGs-based cell type annotation

To enhance cross-species cell type annotation accuracy, we employed the Orthologous Marker Gene Group (OMG) strategy [9]. Orthologous gene groups between apple and four reference species (*Arabidopsis thaliana, Solanum lycopersicum, Oryza sativa*, and *Zea mays*) were identified using OrthoFinder (v2.5.4) [22]. We supplied the top 200 apple cluster-specific marker genes (as determined by Seurat’s FindAllMarkers) as input for ortholog mapping. The apple orthologs identified were then compared against curated marker gene libraries from the reference species. Cell type annotations were inferred based on conserved expression signatures of these orthologous marker groups across species.

### 2.6 XSpeciesSpanner-based cell type annotation

For cross-species cell type annotation using XSpeciesSpanner [14], we provided the complete set of apple protein sequences together with the list of marker genes identified by FindAllMarkers. XSpeciesSpanner generated initial cell identity predictions based on sequence homology and expression profiles. For each cell cluster, the putative cell type with the highest prediction score and a confidence score above 15 was designated as the cluster’s identity.

### 2.7 Cell trajectory analysis

We performed pseudotime trajectory analysis using Slingshot to reconstruct developmental lineages within selected cell populations [23]. For each target cell type, we subset the nuclei from the integrated Seurat object and reprocessed them using a standard workflow (identifying highly variable genes, performing PCA, constructing a shared nearest-neighbor graph, and generating a UMAP embedding). The resulting Seurat objects were converted into SingleCellExperiment format, and the UMAP coordinates were provided to Slingshot as the low-dimensional embedding.

Slingshot was run in an unsupervised mode using cluster labels as input, allowing the algorithm to infer lineage structure and fit smooth principal curves through the transcriptomic space. For each inferred trajectory, Slingshot returned pseudotime values for cells and lineage-specific cell weights, which we used for downstream analyses. To identify genes with dynamic expression along each pseudotime trajectory, we applied tradeSeq [24], fitting generalized additive models with multiple knots to capture non-linear transcriptional changes. Differential expression along the lineages was evaluated using the associationTest, and genes were grouped into stage-specific expression modules based on their smoothed expression patterns. We performed Gene Ontology enrichment analysis for each module to characterize the biological programs associated with distinct phases of lineage progression.

### 2.8 Gene Ontology term enrichment analysis

To construct the Gene Ontology (GO) library for apple, we utilized EGGNOG-MAPPER (v.2.1.12) [25] to predict all proteins. Subsequently, differentially expressed genes (DEGs) from different pseudotime stages were used to perform GO enrichment analysis. GO terms with adjusted P-value < 0.05 were considered significantly enriched. This step was implemented using the enricher function in the clusterProfiler R package [26]. We then visualized the top 10 most significantly enriched terms (with the highest -log_2_(p-value)) from the Molecular Function (MF), Cellular Component (CC), and Biological Process (BP) categories separately in a bubble plot using ggplot2.

### 2.9 Fluorescence imaging of stomata

A first leaf from the plumule was stained with the membrane-specific fluorescent dye FM4-64. Fluorescent signals from abaxial epidermis were captured using a fluorescence microscope (ECLIPSE Ti2-E; NIKON) with 543 ± 11 nm/ 600 ± 26 nm excitation/emission filters.

## 3. Results

### 3.1 Selection of an optimal reference genome improves single-nucleus transcriptome profiling in apple

Young apple seedlings were subjected to nuclei isolation and snRNA-seq using the BGI DNBelab C Series platform (Fig. 1a). The isolated nuclei exhibited high integrity (>90%) and viability (>85%) with minimal debris. After library construction and sequencing, approximately 35,000 high-quality nuclei were obtained, with over 93% passing quality control. Each nucleus generated on average ∼700,000,000 reads and ∼37,000 detected genes (Supplementary Fig. 1 and Supplementary Table 2). These results indicate that our nuclei isolation protocol, coupled with the BGI sequencing workflow, yields reliable, high-quality single-nucleus transcriptomic data for apple. The quality and completeness of the reference genome and gene annotation can strongly influence the accuracy and reliability of RNA-seq data analysis [27]. However, few studies have evaluated this for single-nucleus RNA sequencing (snRNA-seq), especially in non-model crops. We hypothesized that genome assembly quality and accurate gene annotation are key to successful snRNA-seq analysis. To find the best reference for Malus domestica, we compared six public apple genome assemblies (GDDH13, HFTH1, Fuji T2T, Gala, Granny Smith, and GDT2T; Supplementary Table 1). We observed substantial differences in cell abundance and gene capture efficiency across these references. Mapping the same snRNA-seq dataset to each reference genome revealed distinct variations in alignment performance and the average number of genes detected per nucleus. Among the assemblies, GDDH13 consistently performed the best, yielding the highest estimated number of recovered nuclei (35,557) and the greatest mean reads per cell (10,223). It also produced the highest median UMI count (1,939) and median genes per cell (1,233), indicative of superior transcript detection and sequencing sensitivity (Supplementary Fig. 2 and Supplementary Table 2).

**Fig. 1.**
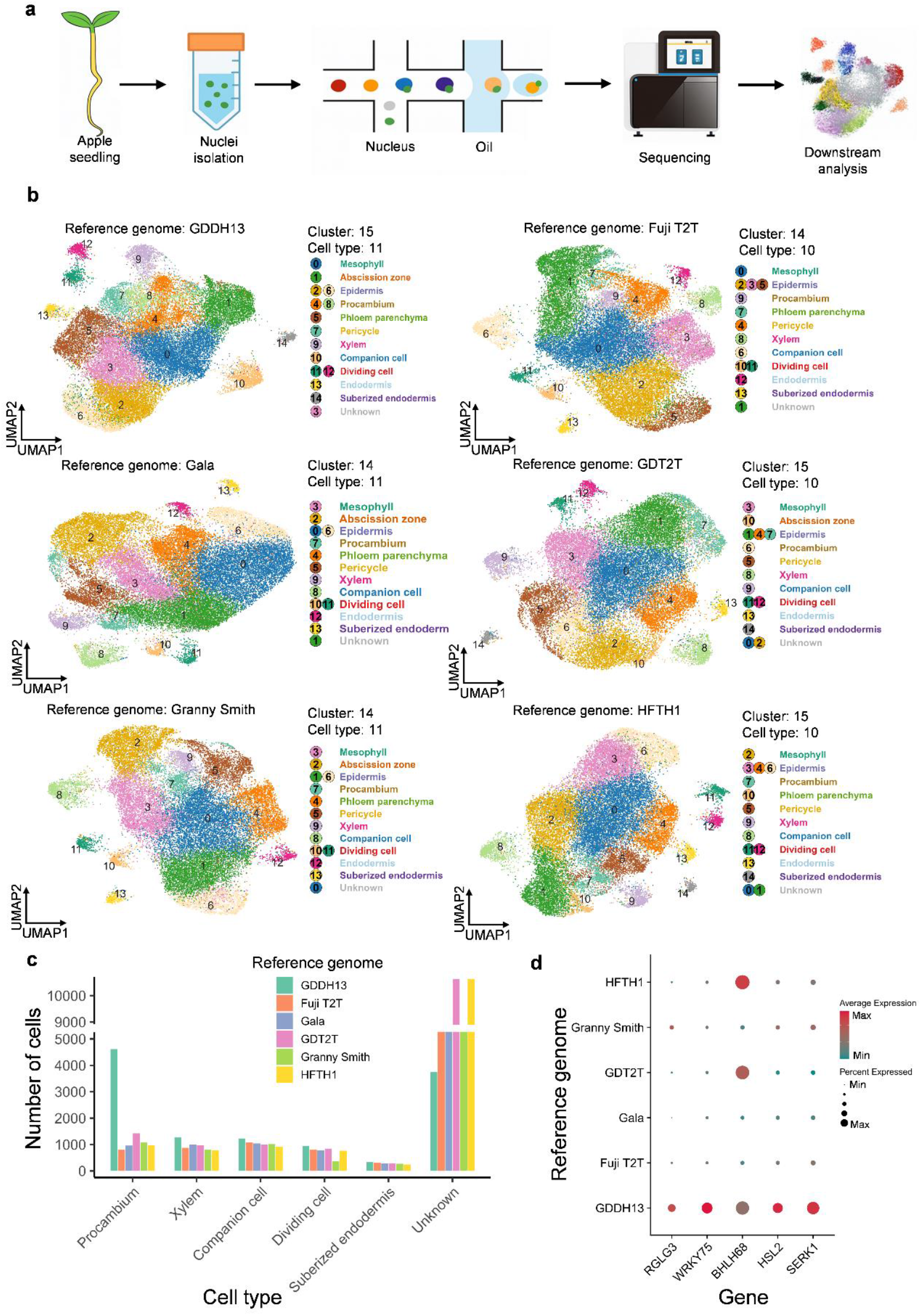
Optimizing the reference genome enhances snRNA-seq profiling in apple. (a) Workflow for single-nucleus RNA sequencing in apple. Nuclei were isolated from young apple seedlings and subjected to high-throughput sequencing and downstream transcriptomic analysis. (b) UMAP visualization of nuclei clustered by transcriptomic profiles based on six different reference genomes: GDDH13, Fuji T2T, GDT2T, Gala, Granny Smith, and HFTH1. Each color indicates a distinct cell type annotated using known marker genes. The number of clusters and identified cell types for each reference genome is indicated. (c) Bar plot showing the number of nuclei assigned to each annotated cell type across the six reference genomes. (d) Dot plot illustrating the expression patterns of representative marker genes across the six reference genomes. Dot size represents the percentage of nuclei expressing the gene, and color intensity reflects the average expression level.

To further assess how the choice of reference genome impacts downstream analysis, we annotated cell types in the apple snRNA-seq dataset using each of the six genome assemblies. We first compiled a set of 29 known plant cell types and their corresponding marker genes from previous reports (Supplementary Table 3). Using homology-based sequence alignment, we identified the apple orthologs of these marker genes in each of the six genomes (Supplementary Table 3), establishing a library for cross-species cell-type annotation and comparative analysis. With the GDDH13 reference genome, the snRNA-seq data yielded the highest transcript counts per cell (exceeding 30,000; Supplementary Fig. 2), resolving 15 transcriptional clusters, of which 14 clusters were confidently annotated as 11 distinct cell types: Mesophyll, Abscission zone, Epidermis, Procambium, Phloem parenchyma, Pericycle, Xylem, Companion cell, Dividing cell, Endodermis, and Suberized endoderm (Fig. 1b). In contrast, the Fuji T2T, Gala, and Granny Smith assemblies each produced only 14 clusters, while the GDT2T and HFTH1 genomes yielded 15 clusters but enabled annotation of only 10 cell types (Fig. 1b). The GDDH13 genome also resulted in the fewest unclassified cells and a greater number of nuclei assigned to well-defined types (such as Xylem and Companion cell), indicating improved annotation precision (Fig. 1c).

To further elucidate why the GDDH13 reference outperformed the others, we examined the expression profiles of key marker genes. Markers such as *RGLG3, WRKY75, BHLH68, HSL2*, and *SERK1* exhibited higher expression levels and stronger cluster specificity when using the GDDH13 genome, which facilitated more accurate cell-type assignments (Fig. 1d). By contrast, the abscission zone marker *HSL2* displayed non-specific expression when reads were aligned to the Fuji T2T or HFTH1 genomes (Fig. 2a), and the phloem parenchyma marker *WRKY75* showed diffuse, weak expression with the GDT2T genome (Fig. 2b), these expression patterns prevented reliable annotation of these cell types. To investigate the underlying cause, we retrieved the *WRKY75* transcript models from the GDDH13, Fuji T2T, and GDT2T genome assemblies. In GDDH13, the *WRKY75* transcript is substantially longer because both the 5′ and 3′ untranslated regions (UTRs) are fully annotated. In contrast, the Fuji T2T and GDT2T annotations lack the longer 3′ UTR of *WRKY75*, which likely reduces the number of snRNA-seq reads that can align to this gene (Fig. 2c). This incomplete gene model annotation may contribute to the lower apparent *WRKY75* expression and subsequent cell-type annotation failures observed with those genomes.

**Fig. 2.**
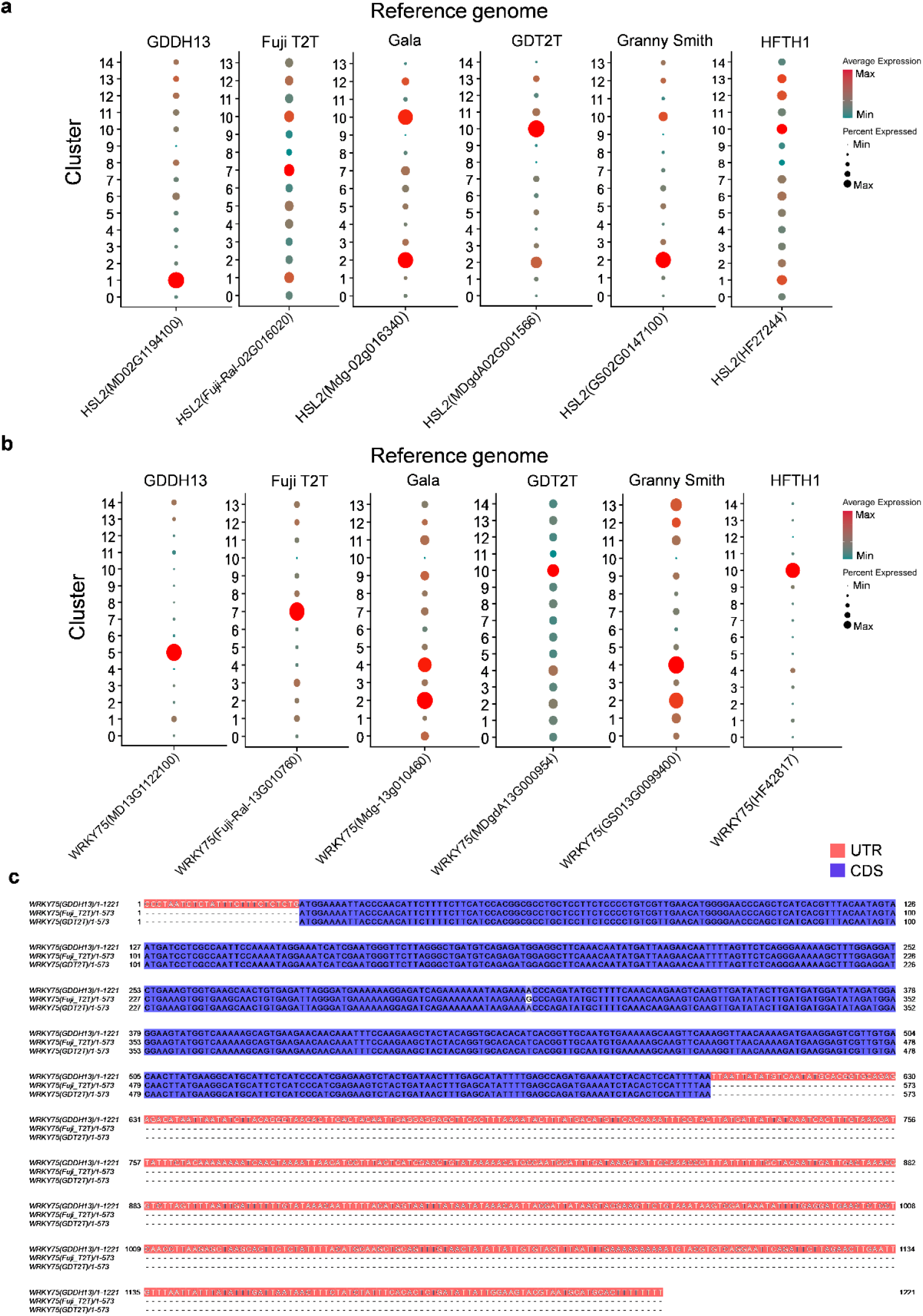
Impact of reference genome choice on *HSL2* and *WRKY75* marker gene expression and transcript structure. (a) Dot plots showing the distribution of *HSL2* expression across different reference genomes (GDDH13, Fuji T2T, Gala, GD T2T, Granny Smith, and HFTH1). Each panel corresponds to one genome, with dot color indicating average normalized expression and dot size indicating the percentage of nuclei expressing the gene in each cluster. (b) As in (a), but for *WRKY75*. (c) Multiple sequence alignment of *WRKY75* transcripts from the GDDH13, Fuji T2T, and GD T2T reference genomes. Untranslated regions (UTRs, red) and coding sequences (CDS, blue) are highlighted, showing the extended 5′ and 3′ UTR annotation of *WRKY75* in the GDDH13 genome.

### 3.2 Cross-validation of cell type annotation using multi-method approaches

Using the GDDH13 assembly and a set of cell type–specific marker genes, we successfully annotated the major cell types in the young apple seedling snRNA-seq dataset (Fig. 3a). For example, clusters 2 and 6 were identified as Epidermis due to their enriched expression of epidermal markers such as *LTPG2, CER6, LTPG1, FDH, PDF1*, and *MYB16*, which are involved in cuticle formation, wax biosynthesis, and epidermal differentiation (Fig. 3a). However, one cluster of cells (cluster 3 in Fig. 3b) remained unassigned and was labeled “Unknown,” underscoring the inherent limitations of relying solely on canonical marker genes for annotation in non-model plants.

**Fig. 3.**
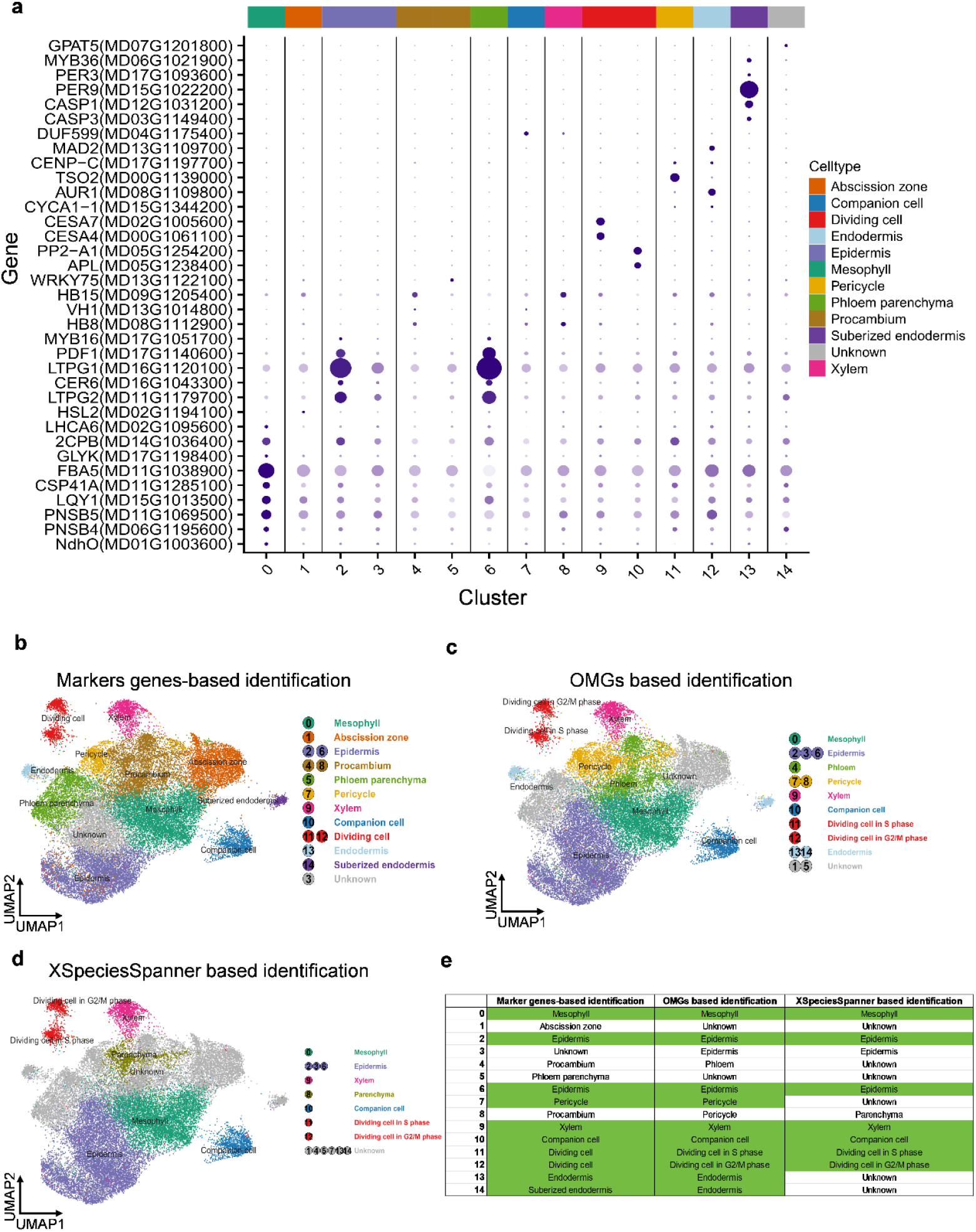
Cell-type annotation of the snRNA-seq dataset using complementary strategies. (a) Dot plot of representative marker genes across clusters (0–14) in the GDDH13 dataset. Dot color shows average scaled expression, and dot size indicates the fraction of nuclei expressing each gene. The color bar above indicates the final cell-type assignment for each cluster. (b) UMAP of nuclei annotated by curated canonical marker genes. (c) UMAP of nuclei annotated using the Orthologous Marker Gene Group (OMG) strategy. (d) UMAP of nuclei annotated using XSpeciesSpanner. (e) Summary table comparing cluster labels across the three methods; green cells indicate concordant cell-type assignments.

Annotation based exclusively on marker genes may be biased if the functions of these genes have diverged among species. Therefore, we incorporated two additional cross-validation approaches—OMGs and XSpeciesSpanner—to improve the robustness and confidence of cell-type assignments. The OMG approach assigns cell identities by detecting conserved groups of orthologous marker genes across species, whereas XSpeciesSpanner compares each query cluster to reference atlases and quantifies their transcriptional similarity. Together, these methods leverage evolutionary conservation and transcriptome-wide pattern matching to support high-confidence cross-species cell-type annotation. In practice, both tools were able to annotate only a subset of the cell types in our apple dataset: OMGs identified 9 cell types, and XSpeciesSpanner identified 7 (Fig. 3c-d), likely reflecting the limited range of cell types represented in their reference databases. Nevertheless, integrating the cross-validation results with our marker gene–based annotations revealed that clusters 0, 2, 6, 7, 9, 10, 11, 12, 13, and 14 had concordant identities across at least two annotation strategies (highlighted in green in Fig. 3e). By contrast, clusters 1 and 5 could not be assigned by either OMGs or XSpeciesSpanner, because their cell types (Abscission zone and Phloem parenchyma) are not present in the reference atlases of those tools (Supplementary Fig. 3-4). Notably, cluster 3 — which was initially unannotated (“Unknown”) based on marker genes alone — was identified as Epidermis by both OMGs and XSpeciesSpanner. This complementary evidence illustrates how a multi-method annotation framework can overcome the limitations of any single approach and substantially enhance the reliability of cell-type identification in non-model plant species.

### 3.3 Branch-specific transcriptional programs reveal divergent differentiation trajectories of vascular tissues

The establishment of a functional vascular system is essential for seedling development, providing long-distance transport, mechanical support, and coordinated tissue growth. To delineate the molecular transitions by which procambial cells differentiate into distinct vascular cell identities, we performed branch-specific pseudotime analyses focusing on the clusters defined as procambium (clusters 4 and 8), phloem parenchyma (cluster 5), and xylem (cluster 9) in Fig. 3b. The resulting pseudotime trajectories revealed that subsets of procambium cells diverged along two well-defined developmental branches, giving rise to a phloem parenchyma lineage (Branch 1) and a xylem lineage (Branch 2). This bifurcation mirrors the canonical developmental pathways of vascular tissue formation in seedlings and provides strong support for the accuracy of our cell-type annotations (Fig. 4a-b).

**Fig. 4.**
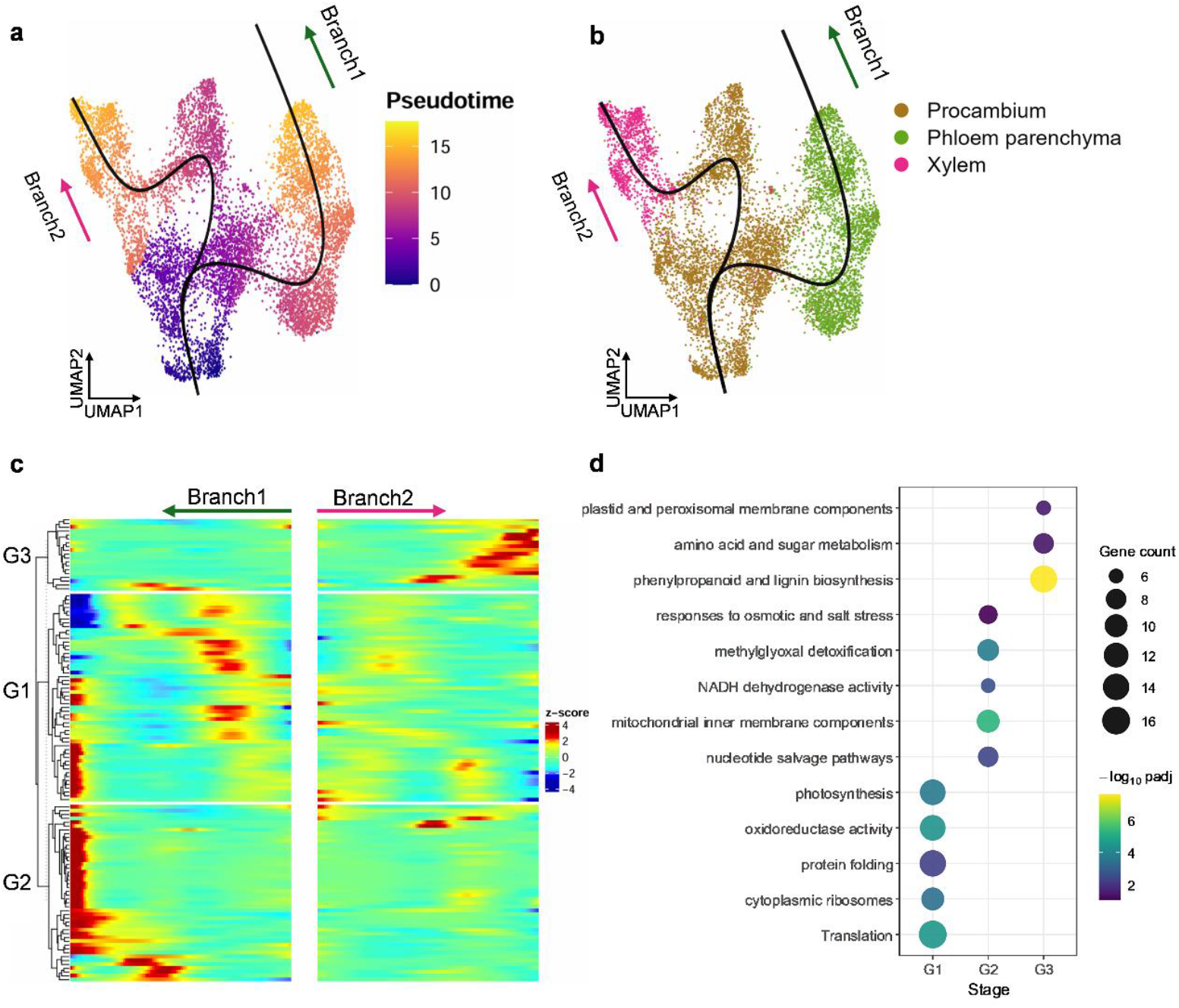
Pseudotime trajectories resolve divergent vascular differentiation. (a) UMAP embedding of procambium, phloem parenchyma, and xylem nuclei from the GDDH13 reference, colored by pseudotime. The fitted principal graph and arrows indicate two developmental branches, with Branch 1 and Branch 2 arising from the procambium. (b) The same UMAP as in (a), colored by annotated cell types, showing the divergence of procambium cells toward phloem parenchyma (Branch 1) and xylem (Branch 2). (c) Heatmap of scaled expression for pseudotime-regulated genes along the two branches. Genes are grouped into three modules (G1–G3) that display distinct expression dynamics along Branch 1 and Branch 2. (d) Gene Ontology enrichment for the three gene modules in (c). The x-axis indicates gene modules (G1–G3) and the y-axis indicates representative enriched biological processes; dot size reflects the number of genes, and color indicates the significance level.

We next analyzed gene expression dynamics along the two differentiation trajectories, with Branch 1 and Branch 2 corresponding to developmental progressions toward phloem parenchyma and xylem, respectively. Genes exhibiting significant temporal variation were grouped into three major expression modules (G1–G3), each representing a distinct transcriptional program activated at a specific stage of vascular differentiation (Fig. 4c). G1 genes, which were strongly upregulated at the early phase of both branches, were enriched for biological processes related to protein translation, ribosome assembly, protein folding, oxidoreductase activity, and photosynthesis (Fig. 4d and Supplementary Fig. 5a). These functions suggest that heightened protein synthesis and metabolic activity are required to initiate procambium differentiation. G2 genes were enriched for pathways in nucleotide salvage, mitochondrial inner membrane organization, NADH dehydrogenase activity, methylglyoxal detoxification, and responses to osmotic and salt stress (Fig. 4d and Supplementary Fig. 5b). These signatures imply that phloem parenchyma maturation (Branch 1) involves enhanced mitochondrial respiration and metabolic adjustments to maintain solute transport and redox homeostasis. G3 genes, predominantly activated along the xylem-directed branch, were enriched for phenylpropanoid and lignin biosynthesis, amino acid and sugar metabolism, and plastid and peroxisomal membrane components (Fig. 4d and Supplementary Fig. 5c). This profile reflects the metabolic reprogramming and secondary cell wall biosynthesis characteristic of xylem differentiation. Collectively, our findings outline a divergent vascular differentiation program comprising an initial shared biosynthetic activation phase followed by two distinct maturation routes—one featuring metabolic and antioxidative specialization in the phloem lineage versus another featuring structural reinforcement and lignin deposition in the xylem lineage. This framework provides a transcriptional basis for the functional divergence of vascular tissues during procambial lineage commitment.

**Fig. 5.**
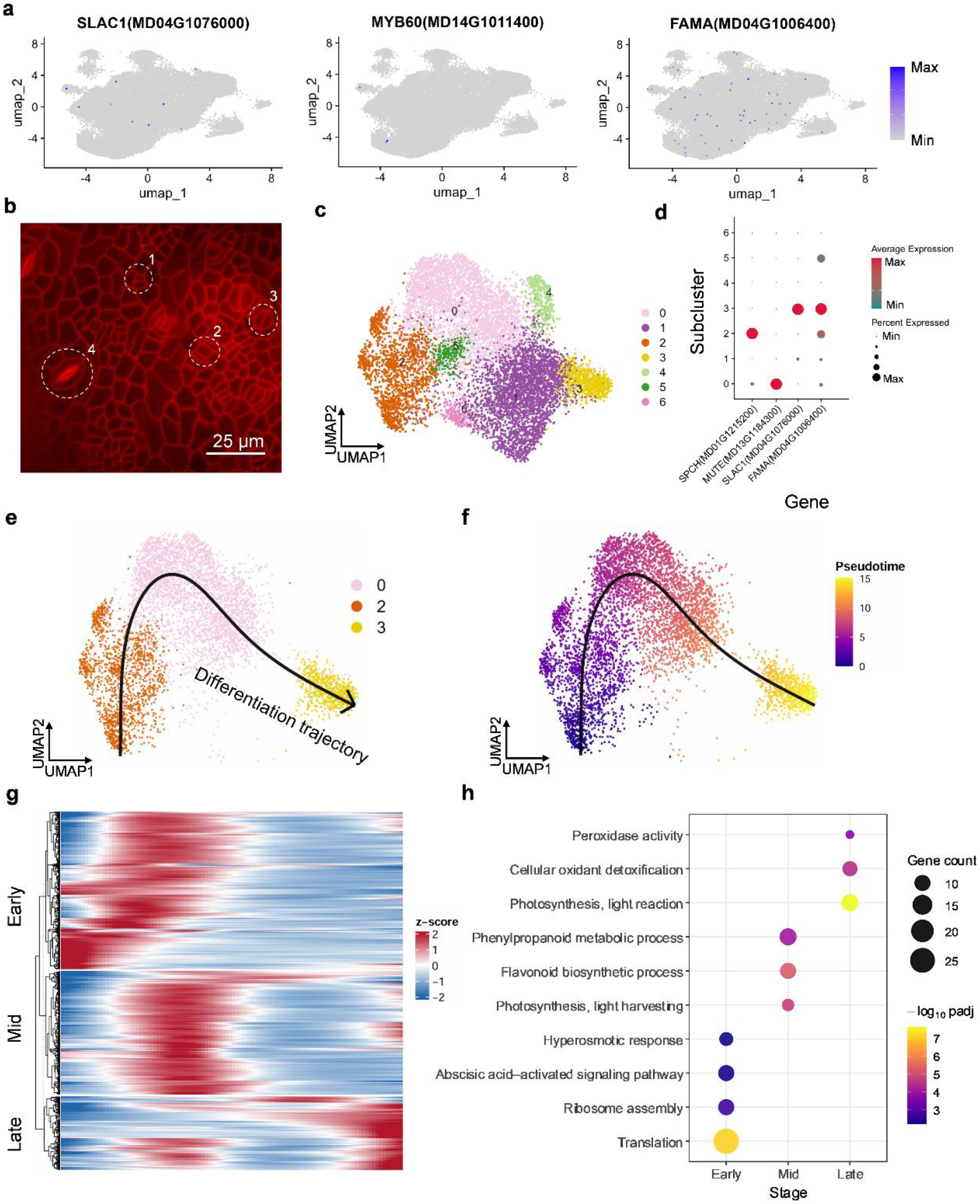
Reconstruction of the stomatal lineage and guard cell differentiation trajectory. (a) Feature plots showing the expression of guard cell regulators *SLAC1, MYB60*, and *FAMA* across all nuclei on the UMAP embedding. (b) Fluorescence image of the apple seedling epidermis. Dashed circles indicate meristemoid, guard mother cell, immature guard cell, and mature guard cell. Scale bar, 25 μm. (c) UMAP visualization of epidermal nuclei (clusters 0, 3, and 6 in Fig. 3b) after subclustering, revealing seven transcriptionally distinct subclusters (0–6). (d) Dot plot showing the expression patterns of key stomatal lineage regulators *SPCH, MUTE, FAMA*, and *SLAC1* across the seven epidermal subclusters. Dot size represents the percentage of nuclei expressing each gene, and color indicates the average expression level. (e) Pseudotime trajectory inferred from subclusters 2, 0, and 3, illustrating the directional progression of stomatal lineage differentiation. Colors denote subclusters. (f) Same trajectory as in (e), colored by pseudotime value, confirming a continuous path from subcluster 2 to subcluster 0 and then to subcluster 3. (g) Heat map of pseudotime-dependent genes along the trajectory in (f). Genes are grouped into three modules corresponding to Early, Mid, and Late developmental phases. (h) Gene Ontology enrichment for the three developmental phases defined in (g). Bubble size indicates the number of enriched genes, and color represents adjusted P values.

### 3.4 Pseudotime reconstruction uncovers molecular transitions during epidermal and guard cell differentiation

The differentiation of guard cells is a critical step in seedling development, as these specialized epidermal cells control gas exchange and transpiration and thus influence the plant’s ability to adjust to environmental conditions. In our snRNA-seq dataset, however, we did not detect a distinct guard cell cluster using classical guard-cell markers, because those markers were expressed at very low levels (Fig. 5a). Consistent with this, microscopic examination of the apple seedling epidermis revealed that stomatal lineage ground cells, guard mother cells, and morphologically immature guard cells were far more abundant than fully developed guard cell pairs (Fig. 5b). This developmental composition suggests that the sampled seedlings were predominantly in an early stage of stomatal-lineage progression, during which mature guard cells are scarce.

To further resolve the early stages of guard cell development, we subclustered the epidermal cells (clusters 0, 3, and 6 in Fig. 3b), which yielded seven transcriptionally distinct epidermal subclusters (0–6; Fig. 5c). Using this finer resolution, we next examined the expression of key regulators in the canonical stomatal lineage pathway (the “*SPCH–MUTE–FAMA*” sequence from asymmetric entry divisions to guard cell differentiation). Consistent with this model, *SPCH* was highly expressed in subcluster 2, *MUTE* was specifically upregulated in subcluster 0, and both *FAMA* and the guard-cell functional gene *SLAC1* were strongly expressed in subcluster 3 (Fig. 5d). Notably, pseudotime analysis of these subclusters confirmed a developmental progression from subcluster 2 to subcluster 0 to subcluster 3, mirroring the temporal order of *SPCH, MUTE*, and *FAMA* activation (Fig. 5e-f). This result demonstrates that our subclustering approach successfully reconstructed the progression of early stomatal lineage differentiation in apple seedlings.

Finally, we profiled gene expression dynamics along the developmental trajectory from subcluster 2 to subcluster 0 to subcluster 3, which revealed three major phases of the stomatal lineage (Fig. 5g). During the early phase, gene expression programs were dominated by pathways related to protein translation, ribosome assembly, and ABA-activated signaling, along with responses to hyperosmotic stress (Fig. 5h and Supplementary Fig. 6a). These features reflect a highly biosynthetic state in which stomatal precursor cells increase protein synthesis and hormonal responsiveness to initiate lineage commitment. The mid phase was marked by a pronounced shift toward flavonoid biosynthesis, phenylpropanoid metabolism, and light-harvesting functions, indicating enhanced plastid activity and secondary metabolic reprogramming (Fig. 5h and Supplementary Fig. 6b). The enrichment of photosynthetic antenna assembly genes suggests that epidermal cells in this intermediate stage begin to acquire light-responsive characteristics. By contrast, the late phase was strongly enriched for genes involved in photosynthetic light reactions, cellular oxidant detoxification, and peroxidase activity (Fig. 5h and Supplementary Fig. 6c), representing signature processes of terminal guard cell specialization that include chloroplast maturation and the activation of ROS-scavenging systems required for functional guard cell formation.

## 4. Discussion

This study presents the first snRNA-seq atlas of young apple seedlings and demonstrates that methodological choices, particularly the selection of the reference genome and the use of multi-faceted cell-type annotation, can critically influence analytical outcomes in non-model plant species. By systematically comparing six apple genome assemblies, we showed that the GDDH13 genome provided markedly superior read alignment quality, higher transcript recovery, and more comprehensive marker gene detection compared with the other apple references (Fig. 1 and Supplementary Table 2). These improvements likely result from GDDH13’s more complete gene models, including fully annotated UTRs that improve read mappability and expression quantification. Our work provides the first systematic evaluation of genome-dependent biases in plant single-nucleus studies. This finding underscores the importance of selecting or developing high-quality genome annotations that are tailored for single-nucleus transcriptomic applications.

Beyond highlighting the advantages of GDDH13 in our evaluation, the results also emphasize that assembly contiguity alone is insufficient to guarantee optimal performance in snRNA-seq analyses of non-model crops. Telomere-to-telomere (T2T) assemblies such as GDT2T and Fuji T2T provide excellent chromosome-level continuity, yet their current structural and functional annotations are less complete than that of GDDH13, exemplified by truncated WRKY75 gene models that lack annotated 3′ UTRs (Fig. 2). Because droplet-based snRNA-seq primarily captures 3′ transcript ends, missing UTR regions decrease the likelihood that reads will be correctly assigned to their genes, which in turn leads to underestimation of key marker gene expression, diminished cluster specificity, and in some cases a failure to resolve entire cell types. These findings are consistent with the broader view that gene model accuracy, UTR completeness, and isoform representation are just as critical as assembly contiguity for single-cell transcriptomic applications, particularly in polyploid or highly duplicated plant genomes [11]. Our study suggests that well-annotated non-T2T references, such as GDDH13, will remain the more reliable choice for single-cell and single-nucleus transcriptomic analyses in apple.

Accurate cell-type annotation is widely recognized as a major bottleneck in plant single-cell research, especially in species lacking curated marker gene resources [7,9]. In our dataset, we found that using only a single annotation approach could yield incomplete or ambiguous classifications, as classical marker genes failed to resolve certain clusters, particularly in rapidly developing cell populations. To overcome this challenge, we integrated three complementary annotation strategies: conventional marker-based labeling, the OMG approach, and XSpeciesSpanner (Fig. 3). In combination, these methods enabled cross-species marker transfer, evolutionary validation, and transcriptome-wide pattern matching. The convergence of evidence from multiple strategies greatly increased our confidence in the cell-type assignments and reduced the risk of misannotation, aligning with recent recommendations advocating multi-method pipelines for non-model species [10,13]. Notably, clusters that could not be resolved by marker genes alone, such as the ambiguous epidermal lineage cluster, were successfully annotated by both OMGs and XSpeciesSpanner, underscoring the importance of integrative frameworks in overcoming species-specific gaps in marker gene knowledge.

Aside from these methodological advances, our dataset provides biologically meaningful insights into early vascular and epidermal differentiation in apple. The pseudotime analyses revealed two bifurcating procambial lineages that give rise to phloem parenchyma and xylem, recapitulating conserved principles of vascular development previously reported in Arabidopsis and poplar [28,29]. The sequential transcriptional changes we identified, beginning with an early increase in translation-related genes and followed by lineage-specific metabolic reprogramming, are consistent with known gene expression programs that control vascular identity and secondary cell wall formation [30] (Fig. 4). Likewise, the reconstructed stomatal lineage in apple closely followed the canonical SPCH– MUTE–FAMA progression described in Arabidopsis [31], suggesting that fundamental regulatory hierarchies of epidermal cell differentiation are broadly conserved in woody perennials. Notably, the predominance of early stomatal-lineage cells in our dataset points to asynchronous epidermal maturation in young seedlings, capturing early stages of guard cell specification that are rarely observed in vegetative tissues (Fig. 5).

## 5. Conclusion

Overall, our study advances both the technical and biological foundations of single-nucleus research in apple. Technically, we demonstrate that reference genome quality and annotation strategy are crucial determinants of analytical accuracy in non-model species and should be explicitly considered in future plant single-cell studies. Biologically, we provide a comprehensive single-nucleus atlas of developing apple seedlings, uncovering both conserved and species-specific features of vascular and epidermal differentiation. This resource lays the groundwork for dissecting gene regulatory networks underlying key developmental processes in fruit trees and may facilitate comparative single-cell studies across woody perennials. Looking ahead, we anticipate that integrating spatial transcriptomics, refined genome annotations, and multi-omics approaches with this atlas will further enhance the resolution and interpretability of developmental programs in apple and related species.

### CRediT authorship contribution statement

**Shupei Hu**: Data curation, Formal analysis, Visualization, Writing – original draft. **Yiqi Liu**: Investigation, Data curation. **Yujie Zhao**: Investigation, Data curation. **Bo Li**: Investigation, Methodology. **Yawen Shen**: Investigation, Methodology. **Yoshiharu Mimata**: Investigation, Visualization. **Kunxi Zhang**: Investigation, Resources. **Pengbo Hao**: Resources, Software. **Jiangli Shi**: Resources, Supervision. **Zhiwei Luo**: Writing – review and editing. **Wenxiu Ye**: Conceptualization, Methodology, Supervision. **Xianbo Zheng**: Conceptualization, Supervision, Funding acquisition. **Jian Jiao**: Conceptualization, Writing – review and editing, Supervision, Project administration.

## Acknowledgments

We thank members of the Grapevine Improvement and Utilization Laboratory at Peking University Institute of Advanced Agricultural Sciences (PKU-IAAS; PI Wenxiu Ye) for helpful discussions and comments on this project. This work was supported by grants from the Key R&D Special Project of Henan Province (251111112900), the Modern Agricultural Industry Technology of Henan Province (HARS-22-09-G3), and Supported by Henan Provincial Science and Technology Research Project (242102110284)

