## Supplementary Figures for "Comparative evaluation of reference genomes and cell-type annotation frameworks for single-nucleus transcriptomic analysis in apple"

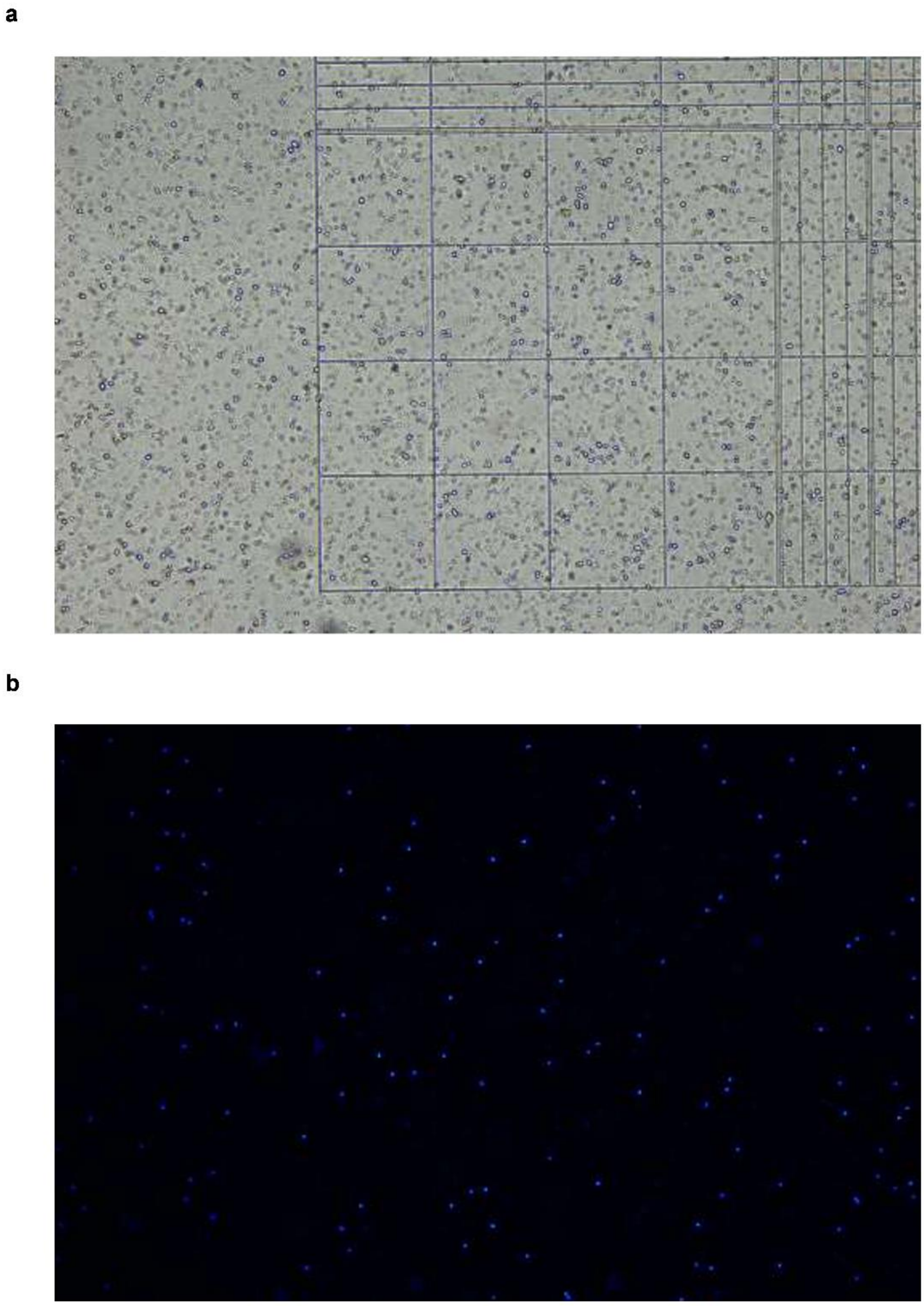


**Supplementary Fig. 1. Microscopic quality assessment of isolated nuclei for snRNA-seq**

(a) Bright-field image of the isolated nuclei suspension on a counting chamber, showing evenly distributed, intact nuclei with minimal debris. (b) Fluorescence image of DAPI-stained nuclei confirming high integrity and purity of the preparation before single-nucleus RNA sequencing.

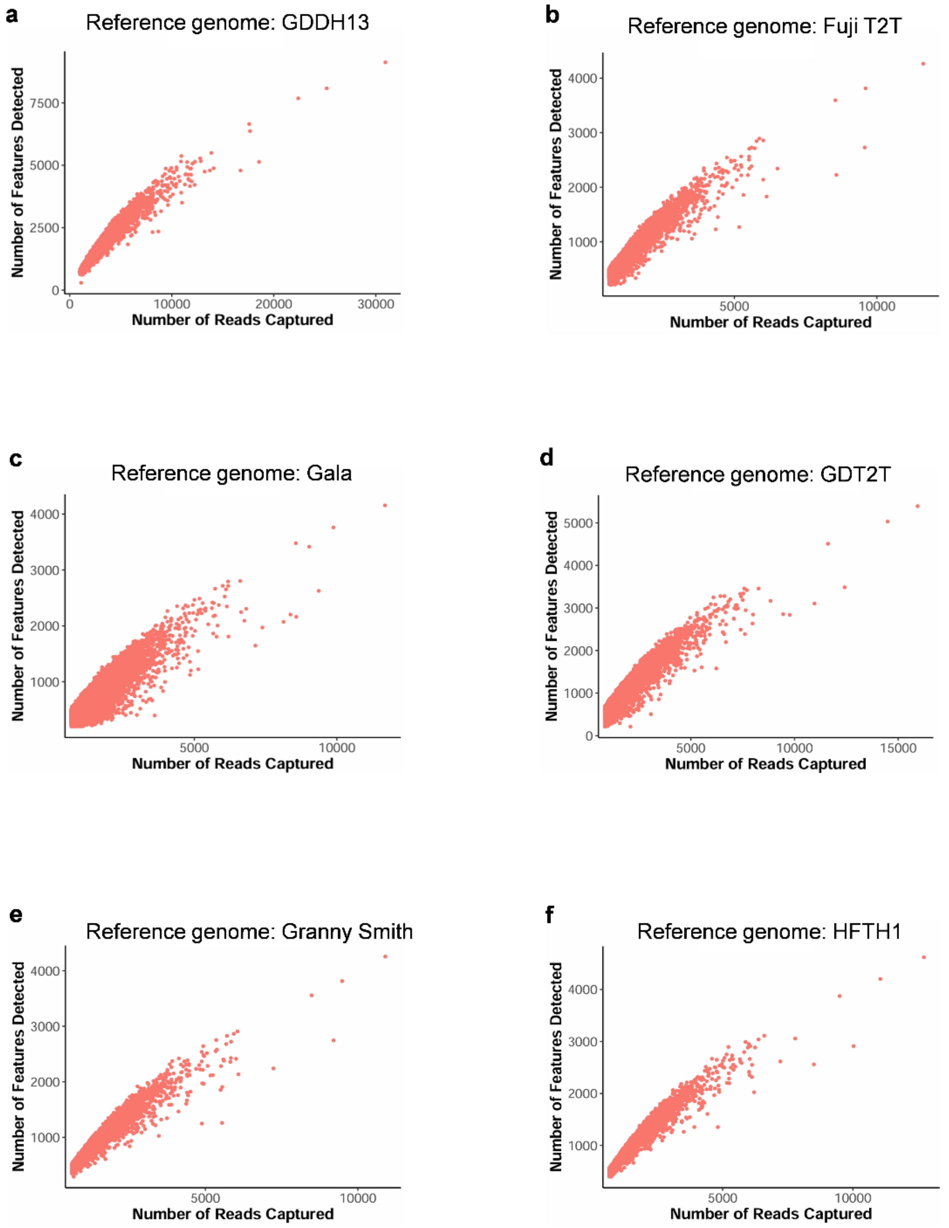


**Supplementary Fig. 2. Quality assessment of snRNA-seq data mapped to different apple reference genomes**

(a–f) Scatter plots showing the relationship between sequencing depth (x-axis, Number of Reads Captured) and transcriptomic complexity (y-axis, Number of Features Detected) for nuclei mapped to the GDDH13, Fuji T2T, Gala, GDT2T, Granny Smith, and HFTH1 reference genomes, respectively. Each dot represents a single nucleus.

a

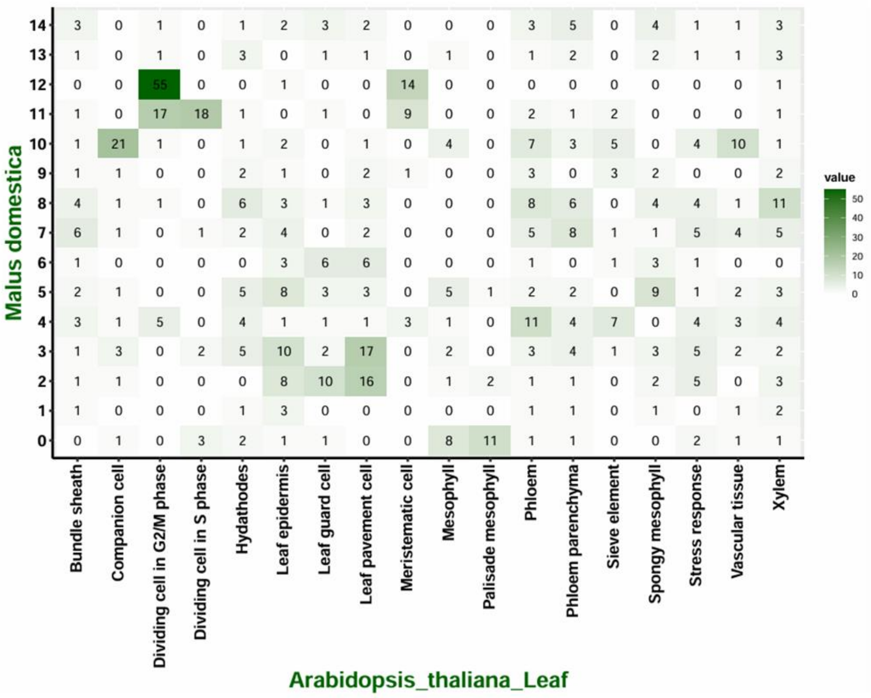
Malus domestica
14
13
12
11
10
9
8
7
6
5
4
3
2
1
0
Bundle sheath
Companion cell
Dividing cell in G2/M phase
Dividing cell in S phase
Hydathodes
Leaf epidermis
Leaf guard cell
Leaf pavement cell
Meristematic cell
Mesophyll
Palisade mesophyll
Phloem
Phloem parenchyma
Sieve element
Spongy mesophyll
Stress response
Vascular tissue
Xylem
Arabidopsis_thaliana_Leaf
value
50
40
30
20
10
0

b

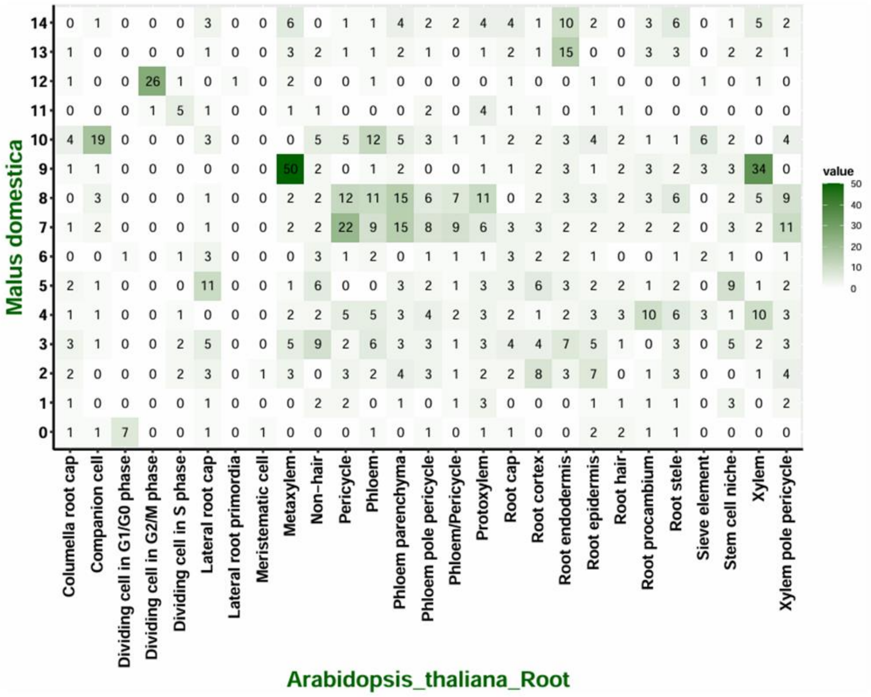
Malus domestica
14
13
12
11
10
9
8
7
6
5
4
3
2
1
0
Columella root cap
Companion cell
Dividing cell in G1/G0 phase
Dividing cell in G2/M phase
Dividing cell in S phase
Lateral root cap
Lateral root primordia
Meristematic cell
Metaxylem
Non−hair
Pericycle
Phloem
Phloem parenchyma
Phloem pole pericycle
Phloem/Pericycle
Protoxylem
Root cap
Root cortex
Root endodermis
Root epidermis
Root hair
Root procambium
Root stele
Sieve element
Stem cell niche
Xylem
Xylem pole pericycle
Arabidopsis_thaliana_Root
value
50
40
30
20
10
0

c

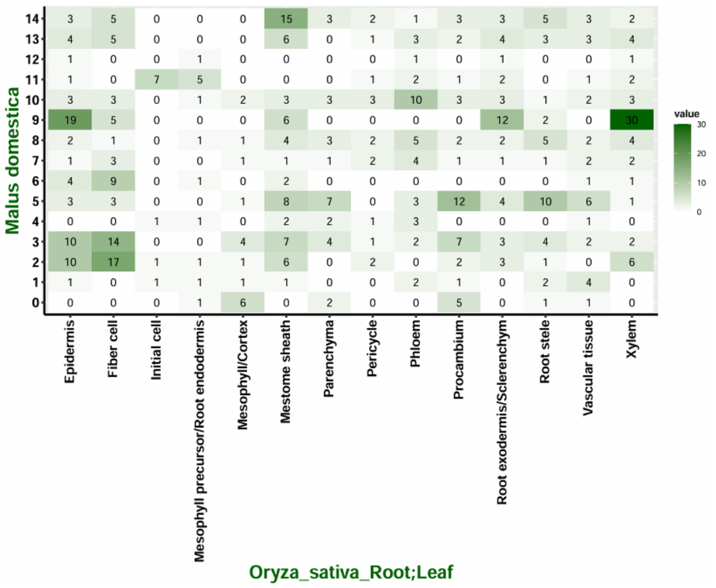
Malus domestica
14
13
12
11
10
9
8
7
6
5
4
3
2
1
0
Epidermis
Fiber cell
Initial cell
Mesophyll precursor/Root endodermis
Mesophyll/Cortex
Mestome sheath
Parenchyma
Pericycle
Phloem
Procambium
Root exodermis/Sclerenchym
Root stele
Vascular tissue
Xylem
Oryza_sativa_Root;Leaf
value
30
20
10
0

d

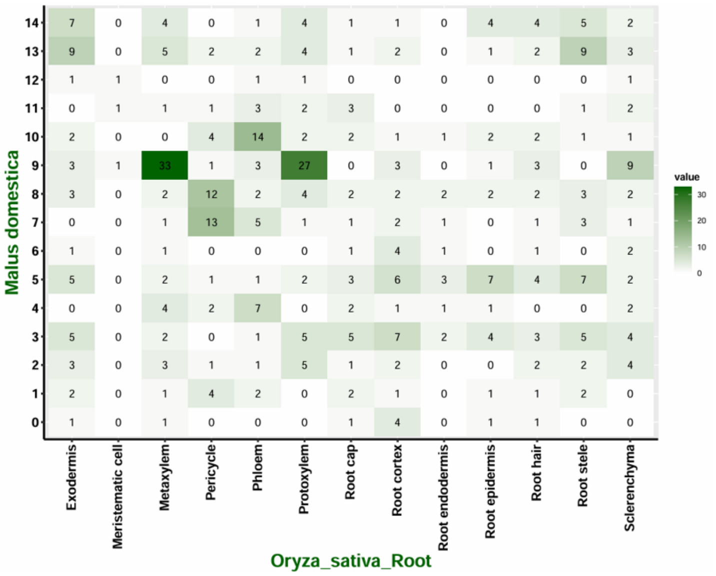
Malus domestica
14
13
12
11
10
9
8
7
6
5
4
3
2
1
0
Exodermis
Meristematic cell
Metaxylem
Pericycle
Phloem
Protoxylem
Root cap
Root cortex
Root endodermis
Root epidermis
Root hair
Root stele
Sclerenchyma
Oryza_sativa_Root
value
30
20
10
0

e

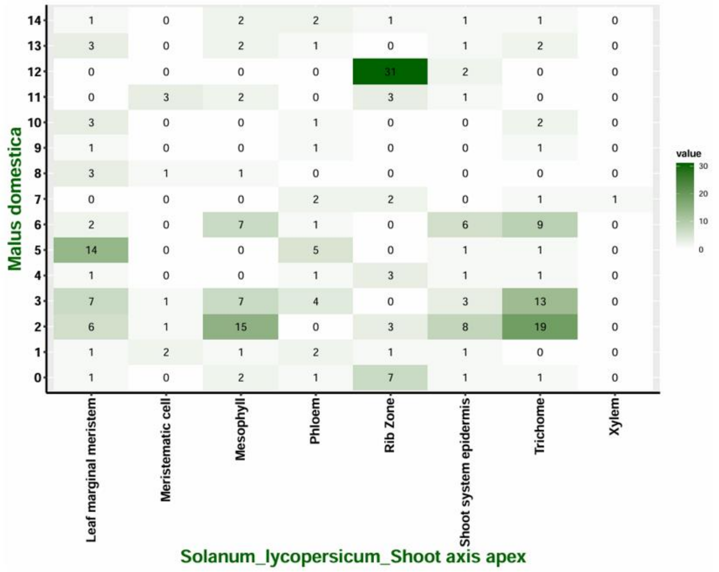
Malus domestica
Solanum_lycopersicum_Shoot axis apex
Leaf marginal meristem
Meristematic cell
Mesophyll
Phloem
Rib Zone
Shoot system epidermis
Trichome
Xylem
value
30
20
10
0

f

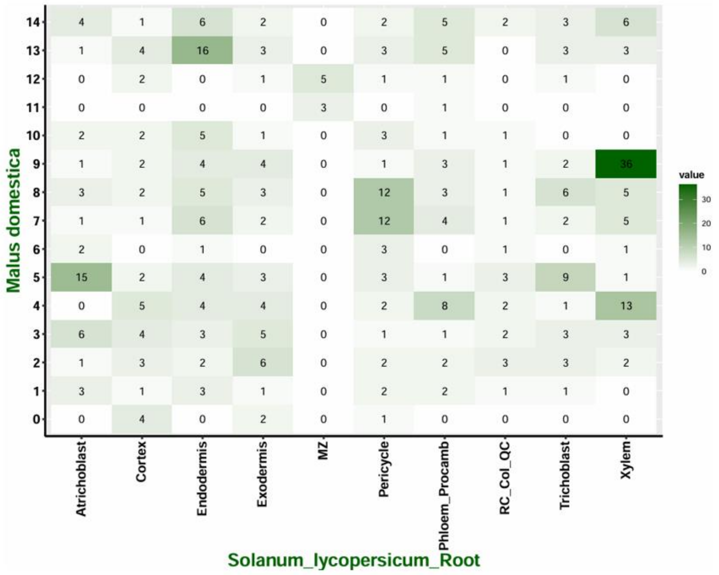
Malus domestica
Solanum_lycopersicum_Root
Atrichoblast
Cortex
Endodermis
Exodermis
MZ
Pericycle
Phloem_Procamb
RC_Col_QC
Trichoblast
Xylem
value
30
20
10
0

g

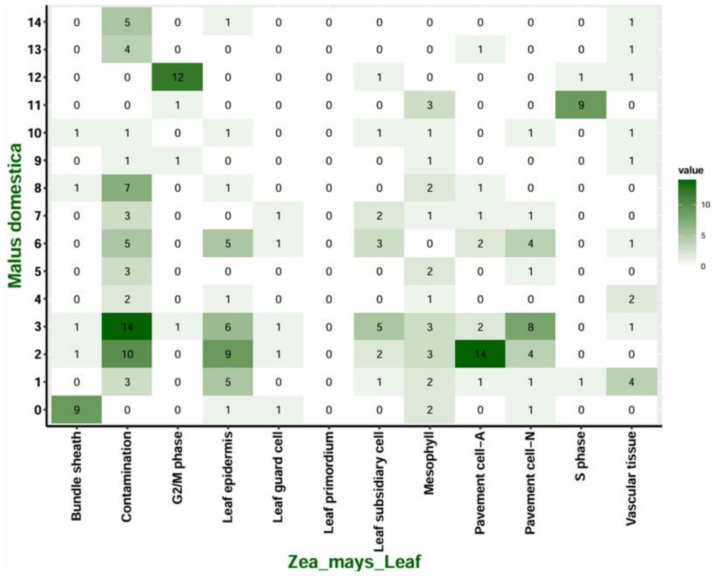


h

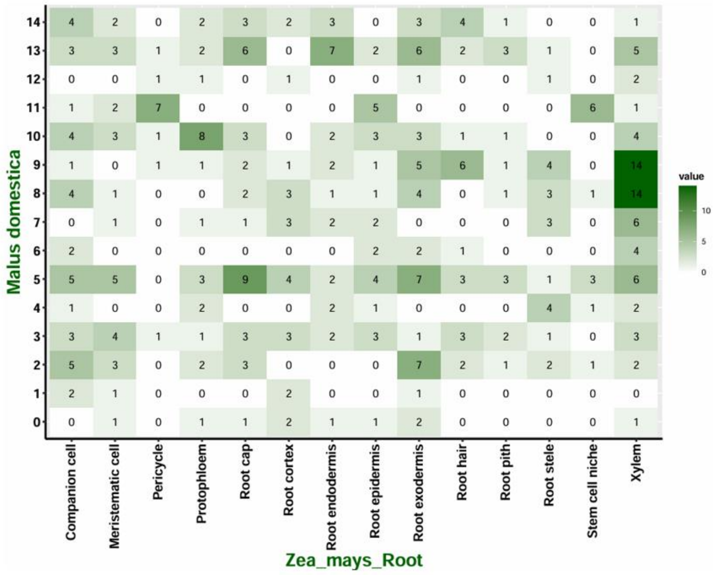


**Supplementary Fig. 3. Cross-species cell-type annotation of the apple young seedling using the Orthologous Marker Gene (OMG) strategy**

(a, b) Heatmaps showing the overlap between Malus domestica cluster marker genes (rows; clusters 0–14 as in Fig. 2b) and cell-type marker sets from *Arabidopsis thaliana* leaf (a) and root (b) single-cell atlases (columns). (c, d) Corresponding OMG-based comparisons with *Oryza sativa* leaf (c) and root (d). (e, f) Comparisons with *Solanum lycopersicum* shoot apical meristem (e) and root (f). (g, h) Comparisons with *Zea mays* leaf (g) and root (h). In all heatmaps, numbers in each tile indicate the count of shared orthologous marker genes between the apple cluster and the indicated reference cell type, and color intensity reflects the magnitude of this overlap.

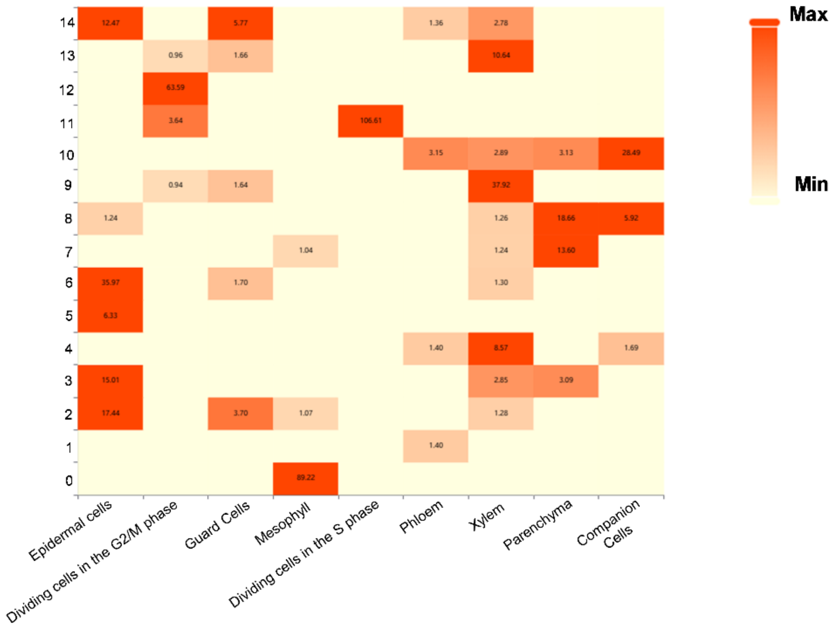


**Supplementary Fig. 4. XSpeciesSpanner-based cross-species cell type annotation of the snRNA-seq**

Heatmap showing XSpeciesSpanner scores for each Seurat cluster (rows, clusters 0–14) across candidate reference cell types (columns). Warmer colors and higher numerical values indicate stronger support for assigning a given cluster to the corresponding cell type.

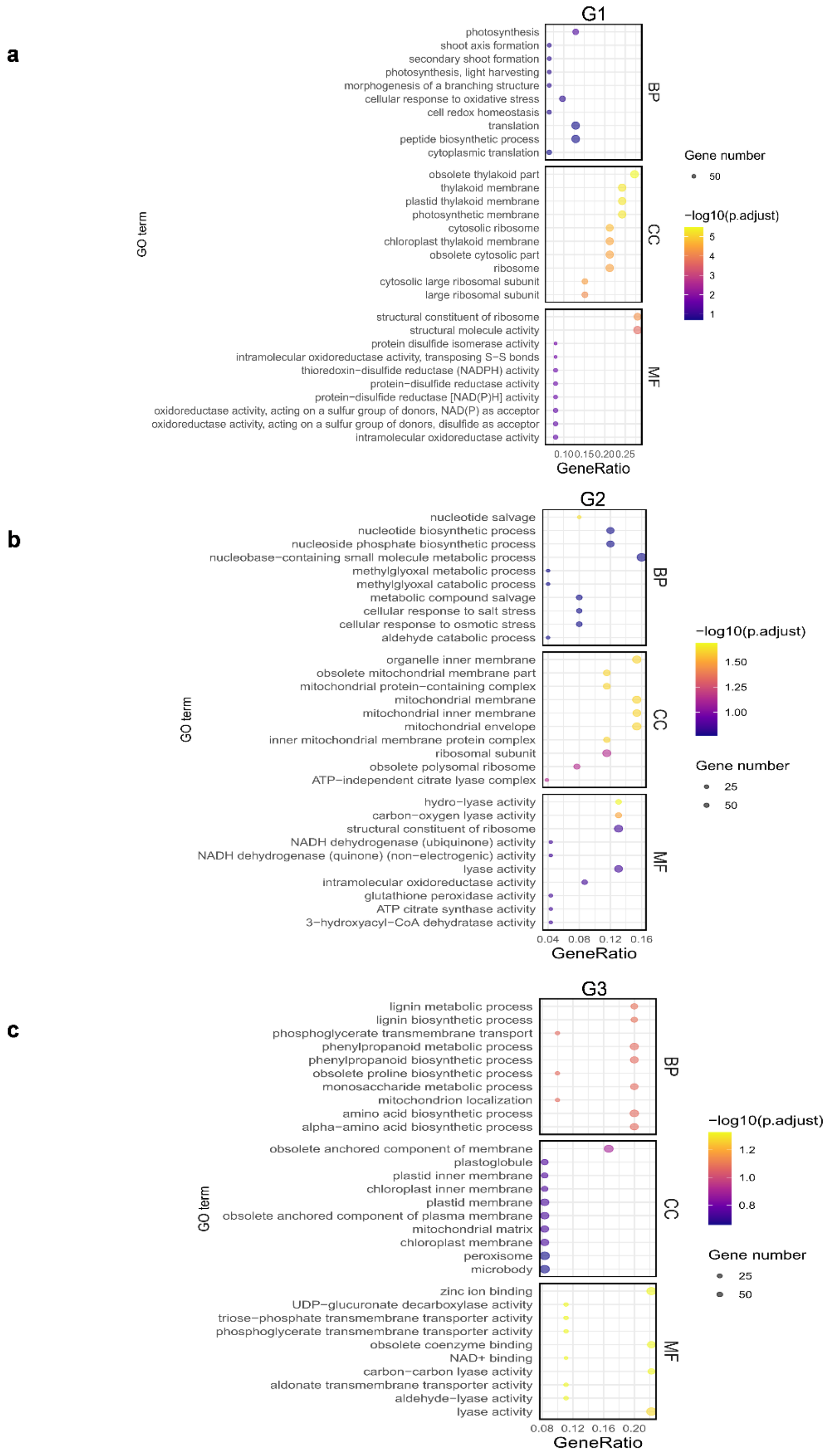


**Supplementary Fig. 5. GO enrichment analysis of gene modules associated with vascular differentiation**

(a–c) Bubble plots showing Gene Ontology enrichment for the three gene modules G1 (a), G2 (b), and G3 (c) identified along the procambium-to-phloem/xylem pseudotime trajectory (Fig. 4c). For each module, significantly enriched biological process (BP), cellular component (CC), and molecular function (MF) terms are displayed. Dot size indicates the number of genes assigned to each GO term, and dot color represents the adjusted P value (–log10 transformed).

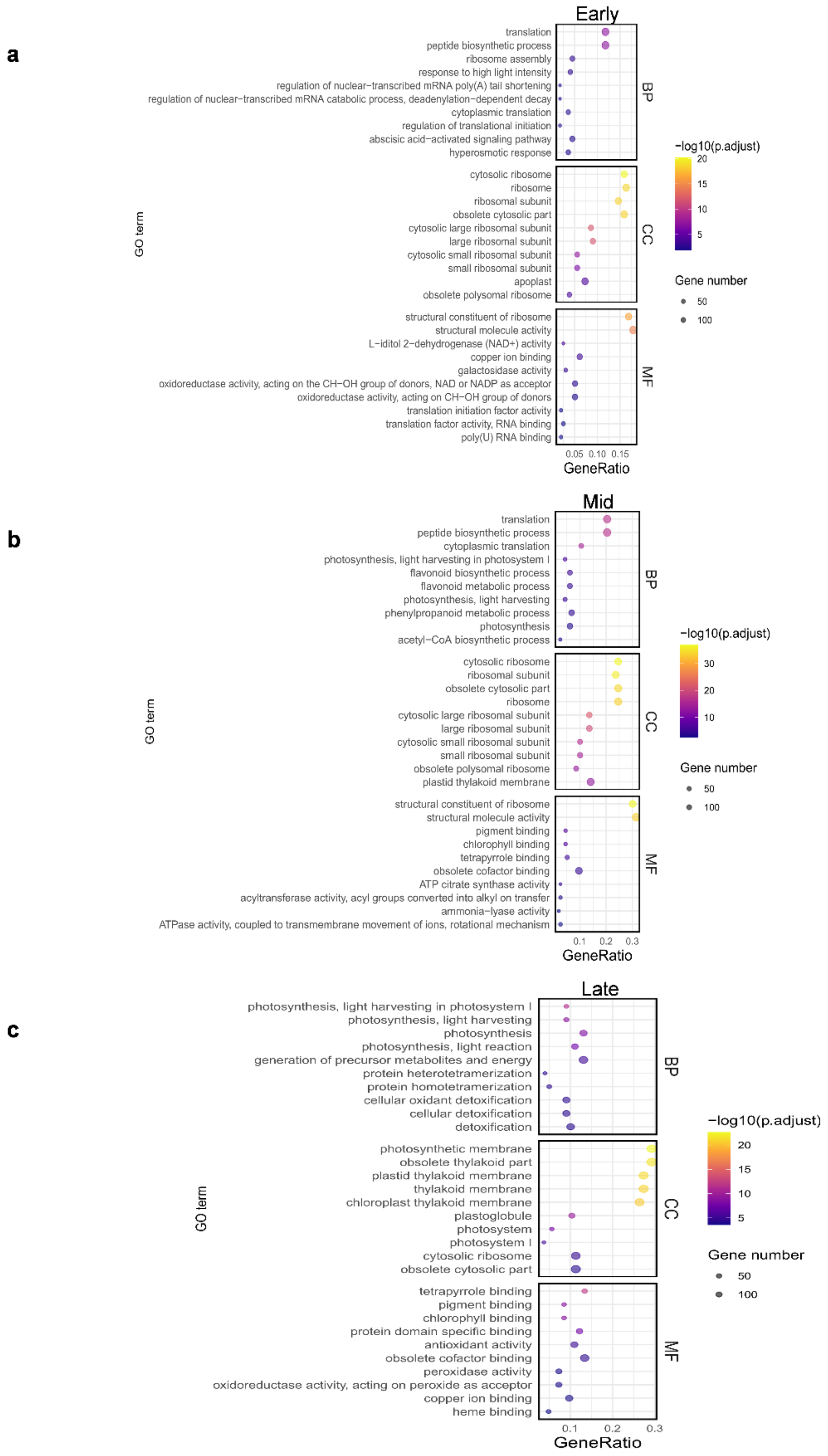


**Supplementary Fig. 6. Gene Ontology enrichment of stage-specific gene modules during stomatal-lineage progression**

(a–c) Dot plots showing GO enrichment for genes preferentially expressed at the Early (a), Mid (b), and Late (c) stages along the guard-cell differentiation trajectory (see Fig. 5g). For each stage, significantly enriched GO terms are grouped into Biological Process (BP), Cellular Component (CC), and Molecular Function (MF) categories. Dot size represents the number of genes associated with each term, and dot color indicates the statistical significance (–log10 adjusted P value). The x-axis denotes the GeneRatio for each enriched GO term.
